# Predicting and Visualizing STK11 Mutation in Lung Adenocarcinoma Histopathology Slides Using Deep Learning

**DOI:** 10.1101/2020.02.20.956557

**Authors:** Runyu Hong, Wenke Liu, David Fenyö

## Abstract

Studies have shown that STK11 mutation plays a critical role in affecting the lung adenocarcinoma (LUAD) tumor immune environment. By training an Inception-Resnet-v2 deep convolutional neural network model, we were able to classify STK11-mutated and wild type LUAD tumor histopathology images with a promising accuracy (per slide AUROC=0.795). Dimensional reduction of the activation maps before the output layer of the test set images revealed that fewer immune cells were accumulated around cancer cells in STK11-mutation cases. Our study demonstrated that deep convolutional network model can automatically identify STK11 mutations based on histopathology slides and confirmed that the immune cell density was the main feature used by the model to distinguish STK11-mutated cases.

## 1. Introduction

Non-small cell lung cancer is the most common type of lung cancer accounting for more than 80% of lung tumor malignancy cases, among which 50% are adenocarcinoma (LUAD) [1]. STK11 is a critical cancer related gene that provides instructions for making a tumor suppressor, serine/threonine kinase 11 [2]. About 24% of all adenocarcinoma cases are STK11-mutated, and molecular studies have shown that STK11-mutation plays an important role in influencing the tumor immune environment including the intratumoral immune cell densities [1]. As a result, many researchers suggested that precision immuno-therapy approaches should take STK11 status of individual tumors into consideration [3–5]. In recent years, deep-learning-based methods have been proved to be able to capture morphological features on tumor images that are associated with molecular features such as mutations, subtypes, and immune infiltration [6–10]. Here, we trained a deep learning model that can determine LUAD patients’ STK11 mutation status based on histopathology slides with high performance. Visualization of the key features learned by the model confirmed that STK11 mutation is associated with the density of immune cells near cancer cells. Practically, this model is capable of providing guidance to immunotherapy in a faster, more convenient, and less expensive way by examining histopathology images without doing sequencing analyses.

## 2. Materials and Methods

Inception-Renet-v2, a modified version of Inception-v4 with residual connection derived from the original InceptionNet, was used as the architecture of the deep learning model for this project [11–13]. Figure 1 shows the general workflow. 541 scanned diagnostic histopathology slides from 478 patients with STK11 mutation status were downloaded from Genomic Data Commons (GDC) of National Cancer Institute (NCI). The data were then separated into training (80%), validation (10%), and testing (10%) sets at per-patient level. Due to the large size of the slides, they were cut into 299-by-299-pixel tiles at 20X magnification level and background was omitted. The model was trained from scratch at per-tile level with batch size of 64 and dropout keep rate of 0.3. The training process stopped when either training or validation loss did not decrease for more than 10000 iterations to avoid overfitting. When training loss reached minimum at some point, a 100-iteration validation was performed. The model was saved as the best performing one only when both training and validation losses were at minimum.

**Figure 1.**
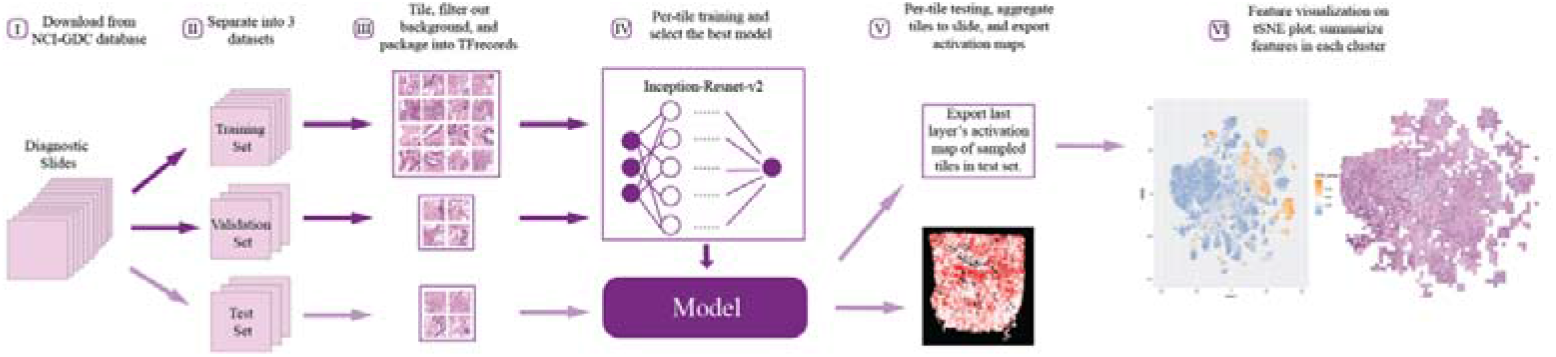
The general workflow of data preprocessing, model training and evaluation, and feature visualization.

## 3. Results

The model achieved per-slide level area under ROC curve of 0.795 (95% CI: 0.601-0.988) and 0.696 (95% CI: 0.692-0.7) at per-tile level (Figure 2). The top-1 accuracy with cutoff at 0.5 was 0.855 (95% CI: 0.742-0.931) at per-slide level and 0.837 (95% CI: 0.835-0.839) at per-tile level. Considering this is a molecular feature prediction task and the labels are at per-slide level only, we believe that these results are quite decent and successful.

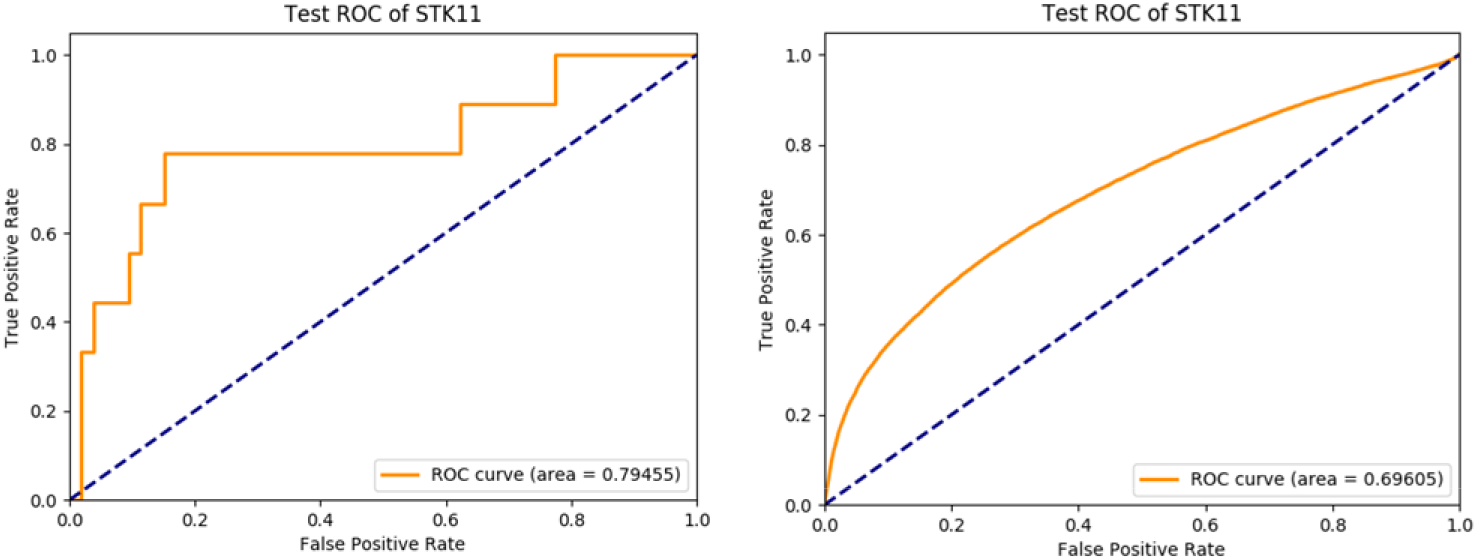
slide level ROC curve and per-tile level ROC curve of the trained InceptionResnetV2 model applying to the test set.

The activation maps before the last fully-connected layer of 30000 randomly selected tiles in test set were recorded. These activation maps were then projected onto a tSNE plot (Figure 3). To have a more straightforward visualization of the features, we put thresholds on prediction scores and randomly selected tiles to represent their corresponding local binned areas on the tSNE space (Figure 4). An experienced pathologist with no previous knowledge in machine learning interpreted patterns in Figure 4 that tiles in the positively predicted clusters (STK11-mutated) generally showing plenty of cancer cells with very few immune cells while a large number of immune cells were present around the cancer cells in the negatively predicted areas (wild type). In addition, most cancer cells were observed in the areas with high positive or negative prediction scores, suggesting that cancer cells were the main focus of the model in making decisions. These findings validated the molecular studies that STK11 mutation decreases the immune response in LUAD patients.

**Figure 3.**
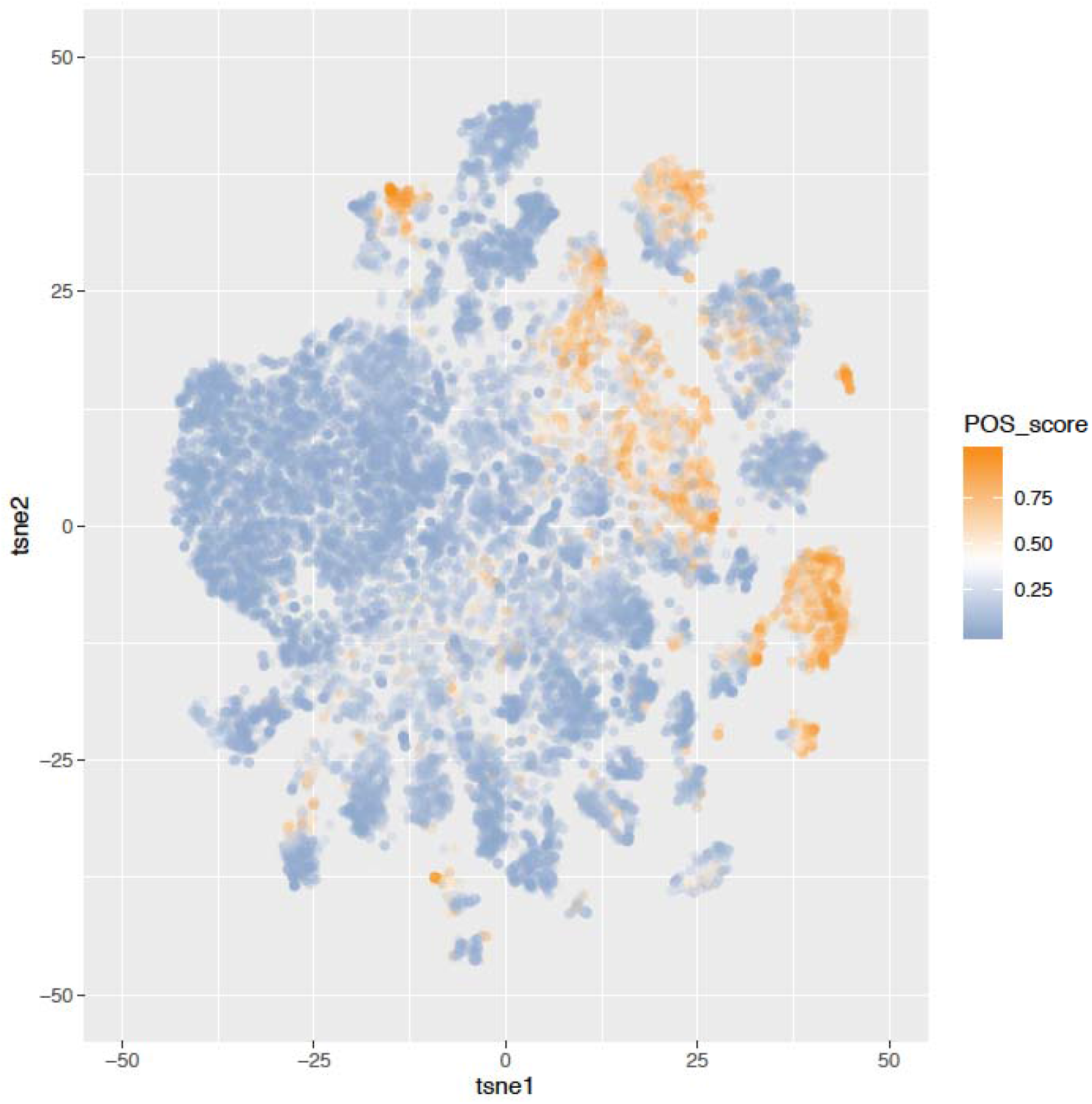
30000 tiles were randomly sampled from the test set. The activation maps before the last fully connected layer of these tiles were represented in the tSNE plot. The color of labels indicating the positive prediction scores of the tiles. Clusters of predicted STK11-mutated and wild type tiles can be observed.

**Figure 4.**
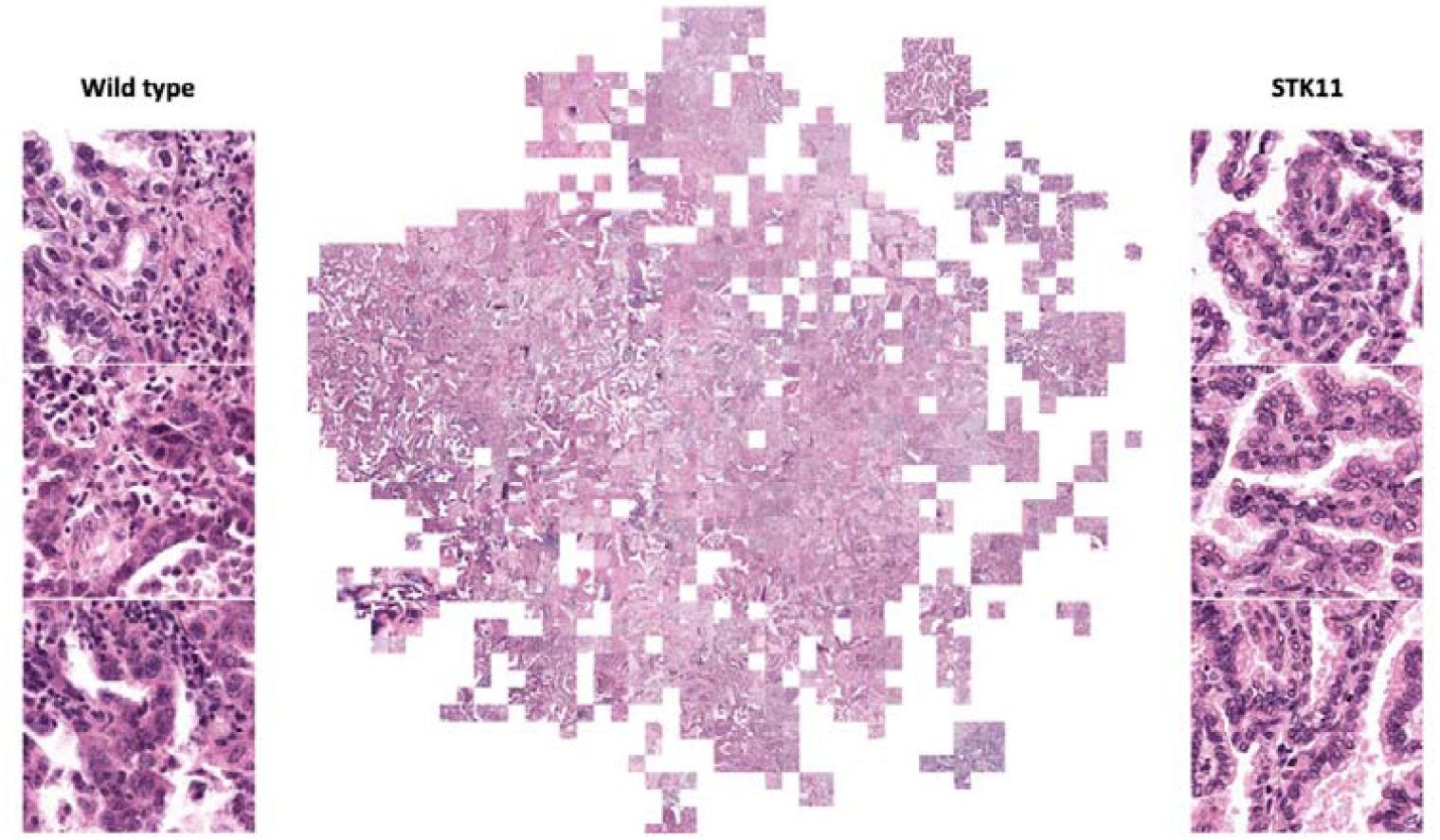
Randomly selected tiles represent binned areas on tSNE space (full resolution figure in supplement). Examples of STK11 mutated and wild type tiles are shown. Cancer cells are the main focuses in these tiles. Predicted STK11 mutated tiles show no immune cells (smaller and darker cells) around cancer cells (larger, lighter, and irregular shape cells) while plenty of immune cells are present in predicted wild type tiles.

## 4. Discussion

The model we trained showed capability in predicting STK11 mutation in LUAD patients based on histopathology images. It has a great potential in providing guidance to immunotherapies in a faster, cheaper, and more convenient way without any sequencing analyses. Scientifically, it confirms the molecular level findings that STK11 mutation leads to less immune response in LUAD tumor from histopathology perspective and links a critical lung cancer molecular feature to a previously unknown morphological pattern. Moving forward, we will continue working on building the connection between cancer molecular features and morphological features using deep learning techniques.

## Supplementary Materials

The following supporting information can be downloaded at: www.mdpi.com/xxx/s1, Figure S1: general workflow

## Author Contributions

Conceptualization, R.H., W.L., and D.F.; methodology, R.H.; software, R.H.; validation, R.H. and W.L.; formal analysis, R.H.; investigation, R.H.; resources, R.H.; data curation, R.H.; writing—original draft preparation, R.H.; writing—review and editing, R.H., W.L., and D.F; visualization, R.H.; supervision, D.F.; project administration, D.F.; funding acquisition, D.F. All authors have read and agreed to the published version of the manuscript.

## Institutional Review Board Statement

Not applicable.

## Informed Consent Statement

Not applicable.

## Data Availability Statement

Genomics data and digital histopathology data can be found at Genomic Data Commons of National Cancer Institute https://gdc.cancer.gov.

## Acknowledgments

We would like to thank the High Performance Computing administration team at NYU Langone Health for maintaining the computational resources of this project.

